# DISA tool: discriminative and informative subspace assessment with categorical and numerical outcomes

**DOI:** 10.1101/2021.12.08.471785

**Authors:** Leonardo Alexandre, Rafael S. Costa, Rui Henriques

## Abstract

**Motivation:** Pattern discovery and subspace clustering play a central role in the biological domain, supporting for instance putative regulatory module discovery from omic data for both descriptive and predictive ends. In the presence of target variables (e.g. phenotypes), regulatory patterns should further satisfy delineate discriminative power properties, well-established in the presence of categorical outcomes, yet largely disregarded for numerical outcomes, such as risk profiles and quantitative phenotypes.

**Results:** DISA (Discriminative and Informative Subspace Assessment), a Python software package, is proposed to assess patterns in the presence of numerical outcomes using well-established measures together with a novel principle able to statistically assess the correlation gain of the subspace against the overall space. Results confirm the possibility to soundly extend discriminative criteria towards numerical outcomes without the drawbacks well-associated with discretization procedures. A case study is provided to show the properties of the proposed method.

**Availability:** DISA is freely available at https://github.com/JupitersMight/DISA under the MIT license.

**Contact:** {} and {}

## 1 Introduction

The discovery of discriminative patterns have proven essential to support predictive and descriptive tasks in biological and medical data domains [2, 4, 7, 10, 11, 13]. In contrast with classic informative patterns, the ability to discriminate an outcome of interest is assessed and, possibly, incorporated in the discovery process [1]. In this context, statistical significance (probability of pattern occurrence against expectations) and discriminative power views are combined in pattern-centric models to aid diagnostics and study regulatory responses to events of interest [1, 7].

Despite the relevance of extending biclustering and pattern mining tasks with discriminative criteria in the presence of target variables, existing contributions are centered on categorical outcomes. To our knowledge, there are no software packages able to robustly assess subspace rules in the presence of numerical outcomes [5, 9]. We propose DISA (Discriminative and Informative Subspace Analysis), a software package in Python (v3.7) to assess patterns with numerical outputs by statistically testing the correlation gain of the subspace against the overall space.

## 2 Methods

A list of subspaces and outcome observations (Figure 2) are the minimum necessary input for DISA. If DISA receives a numerical outcome, a range of values in which samples are valid is determined. DISA accomplishes this by approximating two probability density functions (e.g. Gaussians), one for all the observed targets and the other with targets of the target subspace.

**Fig. 1.**
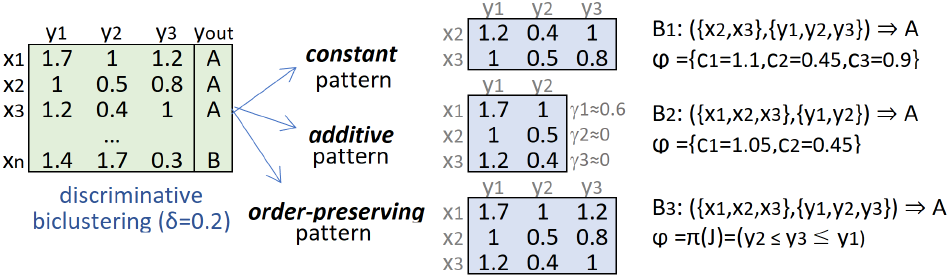
Example of class-conditional subspaces with varying homogeneity. The constant subspace has pattern (value expectations) 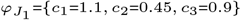, while the order-preserving subspace satisfies the *y*_2_ ≤*y*_3_ ≤*y*_1_ permutation on 3 observations.

**Fig. 2.**
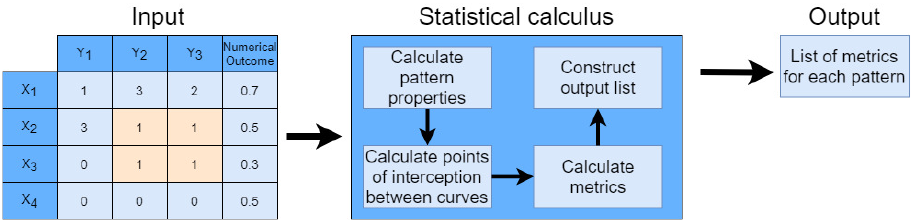
DISA workflow. Input: multivariate data (optional), list of subspaces, outcome variable; Statistical calculus: subspace properties, discriminated ranges from pdf intersection points (numerical outcomes only), and metrics (e.g. statistical significance, gini index, information gain); Output: list of metrics per subspace.

The intersecting points between the two probability density functions is computed to identify the range of values discriminated by the subspace.

DISA supports 53 pattern-centric metrics in total (list in supplementary material) in linear time of observations, returning results in clusters, each ordered by significance to the user.

## 3 Discussion

To illustrate DISA properties, we considered *yeast* dataset [8] and Breast Cancer Wisconsin (diagnostic) dataset [12] (details on preprocessing in supplementary material), both available at the UCI repository [3]. Table 1 provides a synthesis on the DISA assessment over two subspaces produced by BiCPAMS, a state-of-the-art pattern-based biclustering search. P1 corresponds to a subspace in yeast data and P2 to a subspace in breast cancer data (full list is provided in supplementary material).

**Table 1.** Properties of patterns P1, {*mcg*_(4)_, *gvh*_(4)_, *erl*_(0)_, *pox*_(0)_}, and P2, {*radius*_0_(4)_, *perim*_0_(4)_, *area*_0_(4)_, *perim*_2_(4)_, *area*_2_(4)_}, as well as the span of each property and interestingness threshold.

|  | P1 | P2 | Metric interval | Interestingness reference threshold |
| --- | --- | --- | --- | --- |
| Information Gain [13] | 0.05 | 0.02 | [0,1] | $> 0.6$ |
| Gini Index [13] | 0.01 | $8 \times 10^{-3}$ | [0,1] | $> 0.6$ |
| Chi-Squared [2] | **56.10** | 3.3 | $[0, \infty]$ | $> 3.84$ |
| Standardise Lift [10] | **0.73** | **0.71** | [0,1] | $> 0.6$ |
| Stat. Significance [6] | $2 \times 10^{-28}$ | $1 \times 10^{-54}$ | [0,1] | $< 0.05$ |

Figure 3 shows the approximated curves and intersection points associated with P1 and P2 patterns. P1 and P2 approximately discriminate [0.21, 0.5] and [*-*0.17, 0.44] outcome ranges, respectively.

**Fig. 3.**
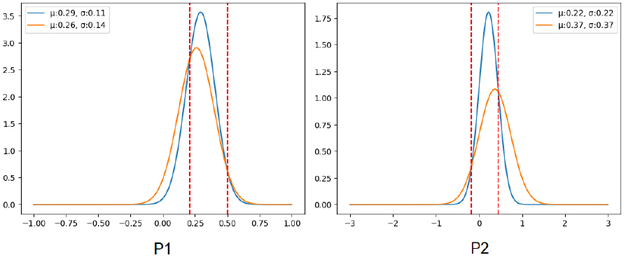
Gaussian intersections between the outcome variable and the subspace of the outcome variable. Blue line represents the Gaussian of the subspace, orange line represents the Gaussian of the original outcome space. P2 intersections, -0.17 and 0.44, correspond to -20 and 55 on the original scale.

## 4 Conclusion

DISA is an open source package capable of robustly assessing the statistical significance and discriminative power of association rules in the presence of numerical and categorical outcomes. DISA implements over 50 metrics, heuristics that can be used to guide the discovery process of discriminative patterns and subspace clusters in biomedical data domains. DISA can be easily embed, therefore aiding the scientific community ability along pattern-centric descriptive and predictive tasks with numerical outcomes.

